# TRIM28 regulates SARS-CoV-2 cell entry by targeting ACE2

**DOI:** 10.1101/2020.08.12.247825

**Authors:** Yinfang Wang, Yingzhe Fan, Yitong Huang, Tao Du, Zongjun Liu, Dekui Huang, Ying Wang, Nanping Wang, Peng Zhang

## Abstract

Severe acute respiratory syndrome coronavirus 2 (SARS-CoV-2) is the cause of coronavirus disease 2019 (COVID-19), it binds to angiotensin-converting enzyme 2 (ACE2) to enter into human cells. The expression level of ACE2 potentially determine the susceptibility and severity of COVID-19, it is thus of importance to understand the regulatory mechanism of ACE2 expression. Tripartite motif containing 28 (TRIM28) is known to be involved in multiple processes including antiviral restriction, endogenous retrovirus latency and immune response, it is recently reported to be co-expressed with SARS-CoV-2 receptor in type II pneumocytes; however, the roles of TRIM28 in ACE2 expression and SARS-CoV-2 cell entry remain unclear. This study showed that knockdown of TRIM28 induces ACE2 expression and increases pseudotyped SARS-CoV-2 cell entry of A549 cells and primary pulmonary alveolar epithelial cells (PAEpiCs). In a co-culture model of NK cells and lung epithelial cells, our results demonstrated that NK cells inhibit TRIM28 and promote ACE2 expression in lung epithelial cells, which was partially reversed by depletion of interleukin-2 and blocking of granzyme B in the co-culture medium. Furthermore, TRIM28 knockdown enhanced interferon-γ (IFN-γ)-induced ACE2 expression through a mechanism involving upregulating IFN-γ receptor 2 (IFNGR2) in both A549 and PAEpiCs. Importantly, the upregulated ACE2 induced by TRIM28 knockdown and co-culture of NK cells was partially reversed by dexamethasone in A549 cells but not PAEpiCs. Our study identified TRIM28 as a novel regulator of ACE2 expression and SARS-CoV-2 cell entry.

## Introduction

Coronavirus disease 19 (COVID-19) is characterized by pneumonia, fever, cough and fatigue, it has rapidly spread to 213 countries and areas, with over 19, 936, 210 total confirmed cases, 732, 499 deaths as of Aug 11, 2020 (WHO, https://covid19.who.int/). COVID-19 is caused by severe acute respiratory syndrome coronavirus 2 (SARS-CoV-2). Recent studies have reported that the entry of SARS-CoV-2 into human cells is mediated by angiotensin-converting enzyme 2 (ACE2) [1], which was previously reported to be the receptor for SARS-CoV [1, 2]. Meanwhile, the entry of SARS-CoV-2 into human cells is facilitated by a host cell serine protease, transmembrane serine protease 2 (TMPRSS2), which cleaves and activates the viral spike glycoporteins to promote virus-cell membrane fusion. The expression levels of ACE2 and TMPRSS2 determine the SARS-CoV-2 cell entry and potentially drive the susceptibility and severity of COVID-19 [3, 4], it is thus of importance to understand the regulatory mechanism of ACE2 and TMPRSS2 expression to develop effective preventive and therapeutic strategies for COVID-19.

Tripartite motif containing 28 (TRIM28) is recently shown to be co-expressed with ACE2 and TMPRSS2 in type II pneumocytes [5], but its roles in the regulation of SARS-CoV-2 receptor and cell entry remain unclear. In this study, we identified TRIM28 as a key factor of ACE2 expression and SARS-CoV-2 cell entry. TRIM28 belongs to TRIM protein family, which comprises of more than 70 members and plays important roles in antiviral restriction, endogenous retrovirus (ERV) latency and immune signaling transduction [6, 7]. For example, TRIM5α restricts human immunodeficiency virus 1 infection by targeting viral capsid protein and causing viral premature [8]. TRIM22 and TRIM32 can induce proteasomal degradation of viral protein, interact with the viral Gag protein and interfere with its trafficking to the plasma membrane [9, 10]. TRIM25 interacts with viral ribonucleoprotein to prevent the viral polymerase from accessing viral RNA template [11]. TRIM28 can associate with viral genome to induce latency of human cytomegalovirus, epstein-barr virus and herpesvirus and transcriptional inhibition of ERVs [12–14]. Moreover, several TRIM proteins have been implicated in autophagy-mediated viral clearance [15]. More detailed TRIM antiviral functions had been well summarized previously [16]. On the other hand, TRIM proteins are shown to have cross-talk with immune system during antiviral restriction. Many TRIM genes are induced following infection with retroviruses, and function as downstream effectors of interferon (IFN)-γ to modulate innate immune responses to viral infections. Knockdown of TRIM genes limits virus-induced interferon production, enhances virus replication and leads to a defect in activations of IFN-regulatory factors (IRFs), nuclear factor-κB and activating protein-1[17]. Furthermore, TRIM genes are crucial regulators of T cell homeostasis, natural killer (NK) and pre-B cell functions [18, 19]. In this study, we investigated the role of TRIM28 in the expression of ACE2 and SARS-CoV-2 cell entry in lung epithelial cells and how TRIM28 contributed to the crosstalk between NK cells and lung epithelial cells. Our results also shed lights on the effect of dexamethasone on TRIM28-modulated ACE2 levels in lung epithelial cells.

## Materials and Methods

### Cell culture and reagents

A549 cells were obtained from ATCC and cultured in RPMI-1640 medium supplemented with 10% fetal bovine serum (FBS, Takara Biomedical Technology Co., Ltd., China), 100 U/ml penicillin and 100 U/ml streptomycin (Invitrogen, Carlsbad, CA, USA) at 37 °C with 5% CO_2_. The NK-92 cells were cultured in the α-MEM medium containing 12.5% FBS, 12.5% heat-inactivated horse serum, 2 mM L-glutamine, 0.1 mM β-mercaptoethanol, 200U/mL of recombinant human interleukin-2, and 100 U/mL of penicillin and streptomycin. Human pulmonary alveolar epithelial cells (PAEpiC), human umbilical vein endothelial cells (HUVECs) and human aortic smooth muscle cells (HSMCs) were purchased from ScienCell (Carlsbad, CA, USA). Human PAEpiCs were cultured in alveolar epithelial cell medium containing epithelial cell growth supplement (ScienCell Company) and 2% FBS in 5% CO2 at 37 °C. Human PAEpiCs at passage 0 were used for all experiments. HUVECs and HSMCs were cultured in EC medium containing EC growth supplement (ScienCell Company) and SMC medium containing SMC growth supplement (ScienCell), respectively. OPTI-MEM medium, lipofectamine RNAi MAX and RNAiso reagent were purchased from Invitrogen.

The cDNA synthesis kits and SYBR Green Master Mix were obtained from Clontech (Mountain View, CA, USA). Dexamethasone and Granzyme B inhibitor Z-AAD-CMKwere purchased from Sigma-Aldrich (St. Louis, MO, USA). Recombinant human IFN-γ was purchased from R&D (Minneapolis, MN, USA). Antibody against TRIM28 was purchased from Santa Cruz (Santa Cruz, CA, USA). Antibodies against phospho-STAT1 and STAT1 and horseradish peroxidase-conjugated goat anti-rabbit antibodies were obtained from Cell Signaling Technology (Beverly, MA, USA).

### Co-culture of NK-92 cells and epithelial cells

NK-92 cells were co-cultured with A549 cells or PAEpiCs at the ratio of 2:1. In the contact culture experiment, A549 cells or PAEpiCs were maintained in 12-well and 24-well culture plates at a density of 1× 10^5^/mL and 2 × 10^4^ /mL, respectively. NK-92 cells were placed directly on top of A549 or PAEpiC cell layer and co-cultured for 24 h at 37 °C. In the non-contact culture experiment, a transwell with 0.4 μm pore polycarbonate membrane insert (Falcon, NJ, USA) was used. The NK-92 cells were cultured in the insert chamber, and the A549 cells were cultured in the bottom chamber.

### SARS-CoV-2 spike-pseudotyped lentivirus and transduction

The SARS-CoV-2 spike-pseudotyped lentivirus that containing expression cassettes for firefly luciferase instead of the VSV-G open reading frame (LV-Spike-nCoV-Luc) [1] was purchased from PackGene Biotech (Guangzhou, China). The target cells were grown in 48-well plates until they reached 50%–60% confluence before they were incubated with pseudotyped virus. At 48 h post-infection, firefly luciferase activities were measured by using Luciferase Reporter Assay System (Promega, Madison, WI, USA) according the manufacturer’s protocol.

### Gene silencing and quantitative PCR (qPCR)

Small-interfering RNAs targeting human TRIM28 (NM_005762.2), ACE2 (NM_021804.3) and IFN-γ receptor 2 (IFNGR2, NM_001329128.1) were synthesized in GenePharma (Shanghai, China) and transfected into cells by using Lipofectamine RNAi MAX. The scrambled siRNA was used as a negative control for siRNA transfection. Total RNA was isolated by using RNAiso reagent. cDNAs were generated by using cDNA synthesis kits (Takara, Japan) according to the manufacturer’s protocol. Subsets of cDNAs were PCR amplified by using SYBR Green Master Mix. The primer and siRNA sequences were shown in the Supplementary Table 1. The level of each transcript was quantified by the threshold cycle method using 18srRNA as an endogenous control.

### Western blotting analysis

For protein expression analysis, the total cellular protein was extracted by RIPA lysis buffer (Thermo Fisher Scientific Inc., Waltham, MA, USA) supplemented with proteinase inhibitor cocktail. Equal amounts of protein extracts were separated by sodium dodecyl sulfate-polyacrylamide gel electrophoresis and transferred on PVDF membranes. Membranes were then immunoblotted with the corresponding primary antibodies. The signals were detected with enhanced chemiluminescence plus (Amersham Pharmacia Biotech)

### Statistical analysis

All the experiments were repeated at least 3 times. The values were expressed as mean ± SEM. Statistical differences were determined to be significant at *P*<0.05. Unpaired Student’s t-test was used to compare difference between two groups. One-way ANOVA and Tukey multiple comparison test were used to calculate significant difference between ≥ 3 groups.

## Results

### Knockdown of TRIM28 increases pseudotyped SARS-CoV-2 cell entry via increasing the expression of ACE2

To study the effects of TRIM28 on ACE2 expression, TRIM28 siRNAs were transfected into A549 cell, a commonly used cell line model for type II pulmonary epithelial cells expressing ACE2 [20, 21]. We observed that TRIM28 knockdown strikingly increased the expression of ACE2 (Figure 1A). TMPRSS2 expression level did not show too much difference between control and TRIM28 knockdown A549 cells (Figure 1B). Meanwhile, knockdown of another TRIM family member TRIM27 did not enhance the expression of ACE2 (Figure 1C). We next employed SARS-CoV-2 spike-pseudotyped virus containing the firefly luciferase gene to evaluate whether TRIM28 controls the cell entry of SARS-CoV-2. As shown in the Figure 1D and 1E, the TRIM28 knockdown cells showed increased luciferase reporter activity, indicating more virus cell entry. Furthermore, ACE2 siRNAs were shown to reverse the increased luciferase reporter activity of TRIM28 knockdown cells, suggesting that ACE2 mediates the increased virus cell entry of TRIM28 knockdown cells. Additionally, we have previously reported the important role of TRIM28 in vascular endothelial cells and smooth muscle cells [22, 23], however, knockdown of TRIM28 did not significantly alter the expression of ACE2 in either of these two types of cells, indicating that expression of ACE2 is regulated by TRIM28 in a cell type-dependent manner (Figure 1F).

**Figure 1.**
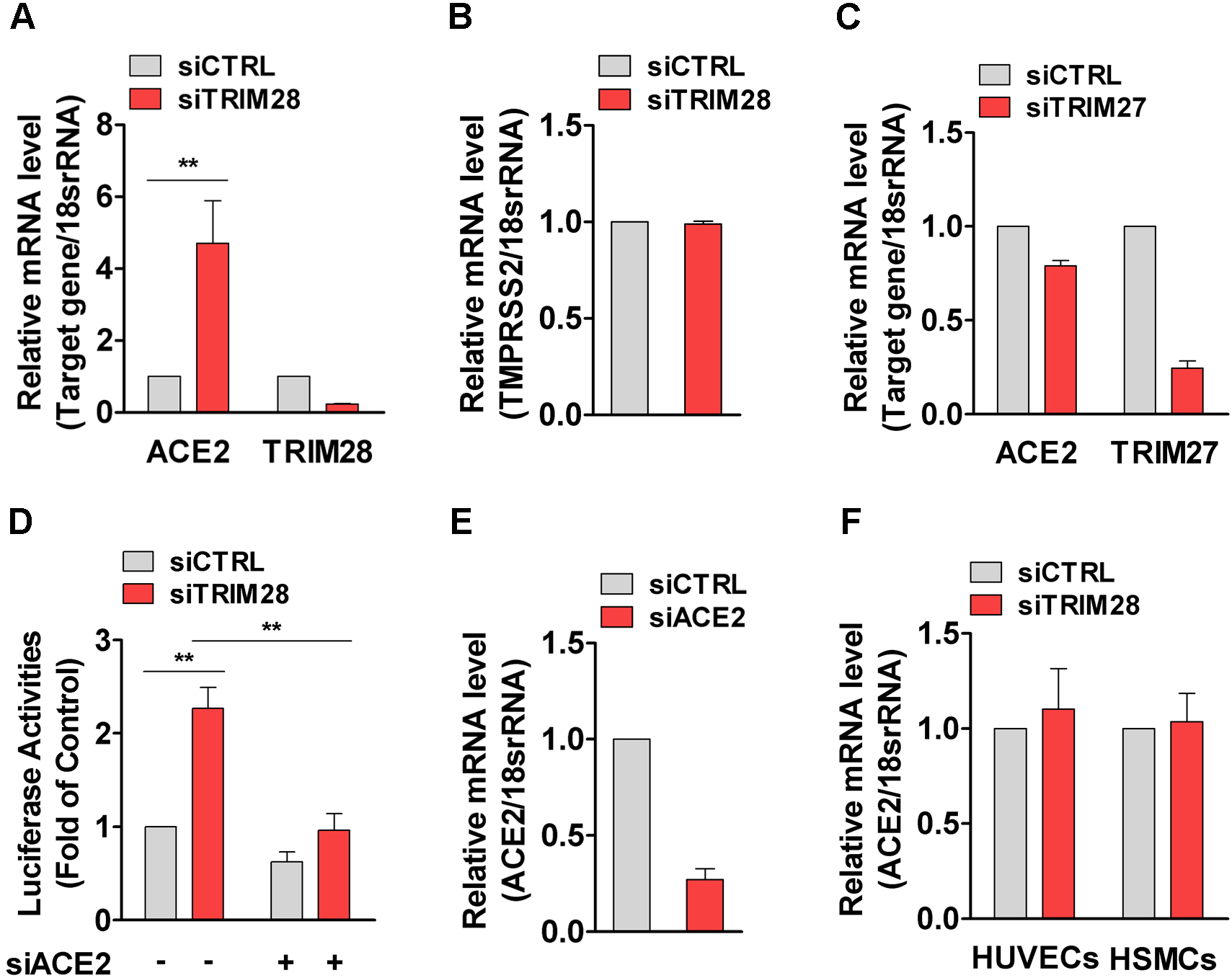
TRIM28 regulates pseudotyped SARS-CoV-2 cell entry in A549 cells. (A-C) The A549 cells were transfected with siRNAs against TRIM28 (A, B) or TRIM27 (C) for 48 h. The mRNA levels of ACE2, TMPRSS2, TRIM28 and TRIM27 were determined by using qPCR method. Data are obtained from 3 independent experiments. \*\**P*<0.01. (D) A549 cells were transfected with TRIM28 siRNA alone or co-transfected with ACE2 siRNA for 24 h, and then infected with LV-Spike-nCoV-Luc for another 48 h before firefly luciferase activities were measured. n = 3. * \**P*<0.01. (E) qPCR assay were used to confirm ACE2 knockdown efficiency. n = 3. (F) HUVECs and HSMCs were transfected with scrambled and TRIM28 siRNA for 48h. The mRNA expression of ACE2 was determined by using qPCR. n = 3.

### Co-culture of NK cells decreases TRIM28 but increases ACE2 expression in A549 cells

The recent discovery that ACE2 is an interferon-stimulated gene in airway epithelial cells has highlighted the important role of defense system in the regulation of ACE2 expression [5]. NK cells can sense, recognize and respond to viral infected cells and regulate the early onset of the inflammatory phase [24–26], and they are recently reported to be present in the bronchoalveolar lavage fluid of patients with severe COVID-19 [27, 28]; however, the role of NK cells in the regulation of TRIM28 and ACE2 remains unclear. We next co-cultured A549 cells with NK-92 cells, which is functional activated NK cells, and observed that the co-culture with NK-92 cells decreased TRIM28 but increased ACE2 expression in A549 cells (Figure 2A-C). Furthermore, NK-92 cells cultured in the IL-2-free medium failed to alter the expression of TRIM28 and ACE2 expression of A549 cells (Figure 2D, E). Interestingly, IL-2 itself did not significantly change TRIM28 expression in A549 cells (data not shown). Since IL-2 maintains the optimal cytotoxic activity of NK cells in culture, our results suggest that the optimal cytotoxic activity of NK cells is required for the regulation of TRIM28 and ACE2.

**Figure 2.**
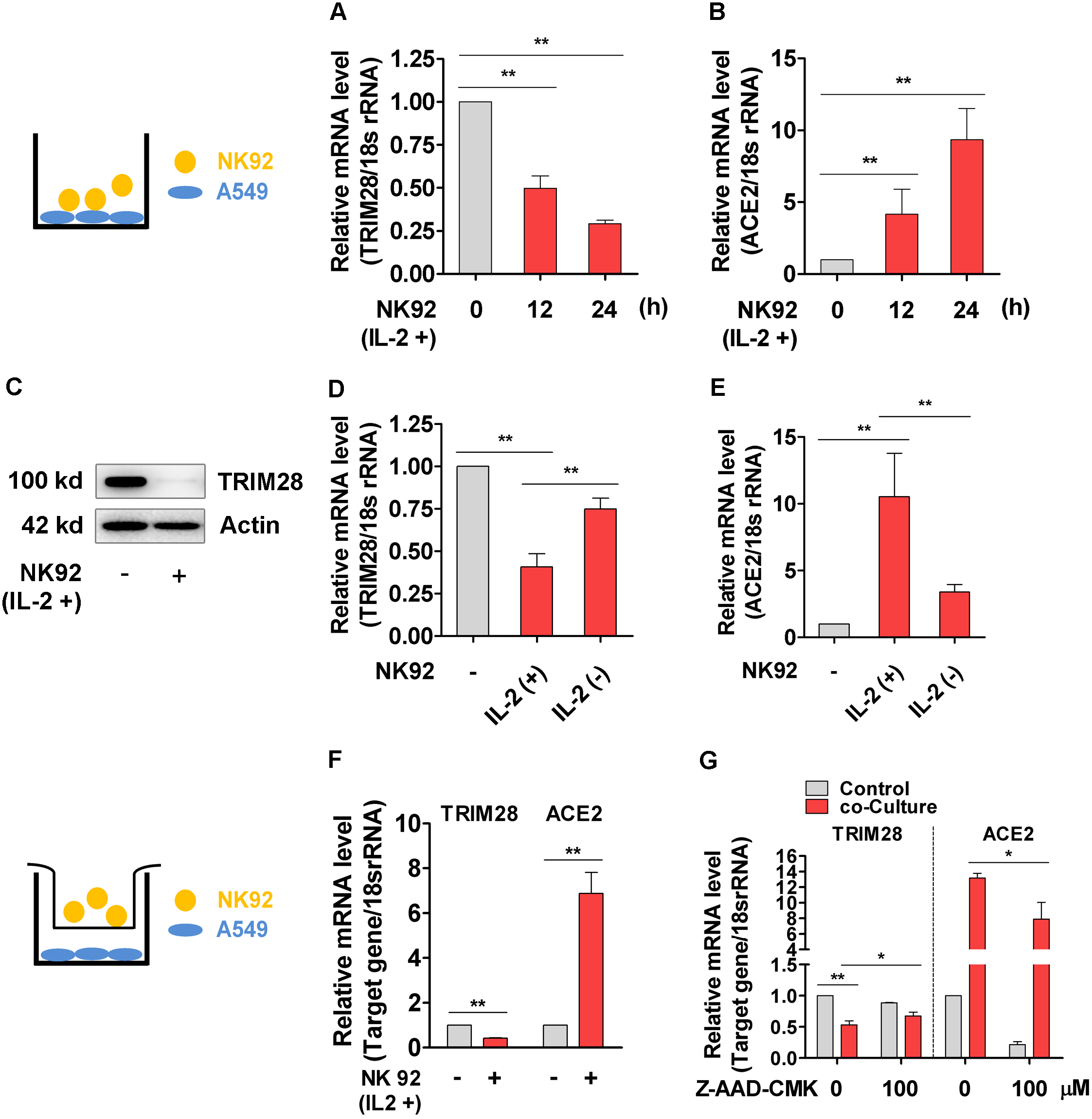
The expression of TRIM28 and ACE2 in A549 cells co-cultured with NK-92 cells. (A-C) The NK-92 cells were placed directly on top of A549 cell layer in 12-well culture plates and co-cultured for 12 or 24 h. The mRNA expressions of TRIM28 (A) and ACE2 (B) were analyzed by qPCR method. n = 3. \*\**P*<0.01. (C) Immunoblot analysis was performed to detect expressions of TRIM28 in A549 cells that co-cultured with NK-92 cells for 24 h. Representative blots were shown. n = 3. (D, E) NK cells were cultured in medium in the presence or absence of IL-2 for 24 h, and then placed on top of A549 cell layer for another 24 h. The mRNA expressions of TRIM28 (D) and ACE2 (E) were analyzed by using qPCR method. n = 3. \*\**P*<0.01. (F) NK-92 cell and A549 cells were co-cultured by placing the NK-92 culture inserts on top of the A549 cell layer in transwell culture plates for 24 h. qPCR assay was used to check mRNA expressions of TRIM28 and ACE2 in A549 cells. n = 3. \*\**P*<0.01. (G) The NK-92 cells were placed directly on top of A549 cell layer and co-cultured in medium containing granzyme B inhibitor Z-AAD-CMK (100 μM) for 24 h. TRIM28 and ACE2 expressions were determined by qPCR method. n = 3. \*\**P*<0.01. \**P*<0.05.

To further determine whether the regulatory effect of NK cells on the expression of ACE2 and TRIM28 of A549 cells is contact-dependent, we co-cultured A549 and NK-92 cells using a transwell co-culture system. We observed that NK-92 cells showed a similar effect on A549 cells that the expression of TRIM28 was significantly decreased whereas ACE2 expression was increased in the transwell co-cutlure system (Figure 2F). These data indicated that paracrine factors from NK cells mediate the regulatory effect of NK cells on the expression of TRIM28 and ACE2 in A549 cells. NK cells are known to defense against viral infections through secretion of IFN-γ, TNF-α, granzyme, GM-CSF and interleukins. We observed that TNF-α (data not shown) and IFN-γ (Figure 4B) did not affect TRIM28 expression in A549 cells. However, Z-AAD-CMK, a granzyme B inhibitor, can partly reverse the inhibition and promotion effects of NK-92 cells on TRIM28 and ACE2 expression of A549 cells, respectively (Figure 2G).

### TRIM28 knockdown- and NK cell-induced ACE2 upregulationin in A549 cells is reversed by dexamethasone

Recent study showed that dexamethasone, a corticosteroid with immunosuppressive and anti-inflammatory effects, can reduce mortality among COVID-19 patients who received mechanical ventilation or oxygen alone [29], which prompted us to evaluate whether dexamethasone can reverse the upregulation of ACE2 and increased pseudotyped SARS-CoV-2 cell entry of TRIM28 knockdown cells. Our results show that dexamethasone reduced ACE2 levels in both basal and TRIM28 knockdown A549 cells (Figure 3A, B). Accordingly, pseudotyped SARS-CoV-2 cell entry was blocked by dexamethasone (Figure 3C). Notably, although dexamethasone did not affect the expression of TRIM28 of A549 cells, it partially reversed the inhibitory effect of NK-92 cell on TRIM28 expression of A549 cells in the co-culture model (Figure 3D). These data indicate that dexamethasone can reverse the expression of ACE2 in A549 cells which were induced by TRIM28 knockdown and co-culture with NK cells.

**Figure 3.**
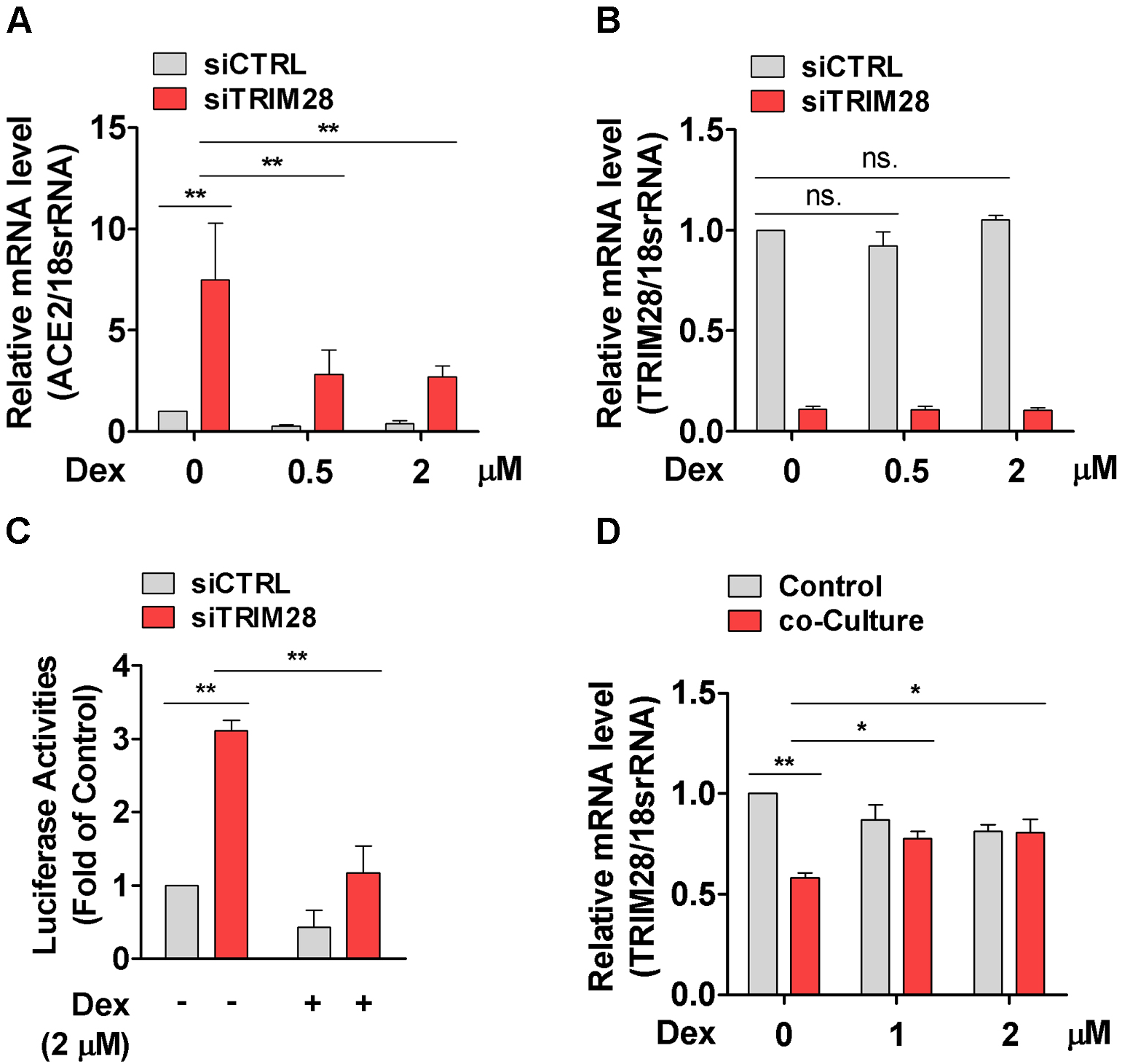
Effect of dexamethasone on TRIM28 knockdown induced ACE2 expression in A549 cells. (A, B) A549 cells were transfected with scramble or TRIM28 siRNA for 32 h and subsequently incubated with dexamethasone for another 16 h. ACE2 (A) and TRIM28 (B) transcripts were analyzed by using qPCR assay. n = 3. \*\**P*<0.01. ns. no significance. (C) Scramble or TRIM28 siRNA transfected A549 cells were treated with dexamethasone for 16 h, and then infected with LV-Spike-nCoV-Luc for another 48 h.The firefly luciferase activities were measured. n = 3. \*\**P*<0.01. (D) The NK-92 cells were placed directly on top of A549 cell layer and co-cultured in dexamethasone containing medium for 24 h. The expression of TRIM28 was determined by qPCR method. n = 3. \*\**P*<0.01. \**P*<0.05.

### TRIM28 knockdown potentiates IFN-γ-induced ACE2 expression in A549 cells

Previous studies have reported that TRIM28 acts as a key mediator of IFN [30], it is recently shown to induce ACE2 expression in lung epithelial cells [5]. We then examined whether TRIM28 affects the IFN-induced upregulation of ACE2 in lung epithelial cells. Our results showed that IFN-γ can up-regulate ACE2 expression in A549 cells (Figure 4A). Importantly, TRIM28 knockdown dramatically potentiated IFN-γ induced ACE2 expression (Figure 4A, B), suggesting endogenous TRIM28 negatively modulates IFN-induced ACE2. We previously observed that TRIM28 can determine expressions of several membrane receptors in vascular cells [22, 23]. Thus, to further identify the molecular mechanism through which TRIM28 controls IFN signaling pathway, we examined the effect of TRIM28 on the expression of receptors of INF-γ, including IFNGR1 and IFNGR2. Our results show that TRIM28 knockdown can increase IFNGR2 but not IFNGR1 expression of A549 cells (Figure 4C). Accordingly, knockdown of TRIM28 strikingly enhanced the phosphorylation of STAT1, the major downstream signaling components of IFN-γ, at both basal level as well as upon IFN-γ stimulation (Figure 4D). Furthermore, our results demonstrated that knockdown of IFNGR2 abolished inductive effect of IFN-γ on ACE2 expression in A549 cells, supporting that INFGR2 is required for IFN-γ to upregulate ACE2 expression (Figure 4E). Taken together, our results suggest that TRIM28 knockdown can potentiate IFN-γ-induced ACE2 expression at least partly through a mechanism involving upregulating IFNGR2 expression in A549 cells.

**Figure 4.**
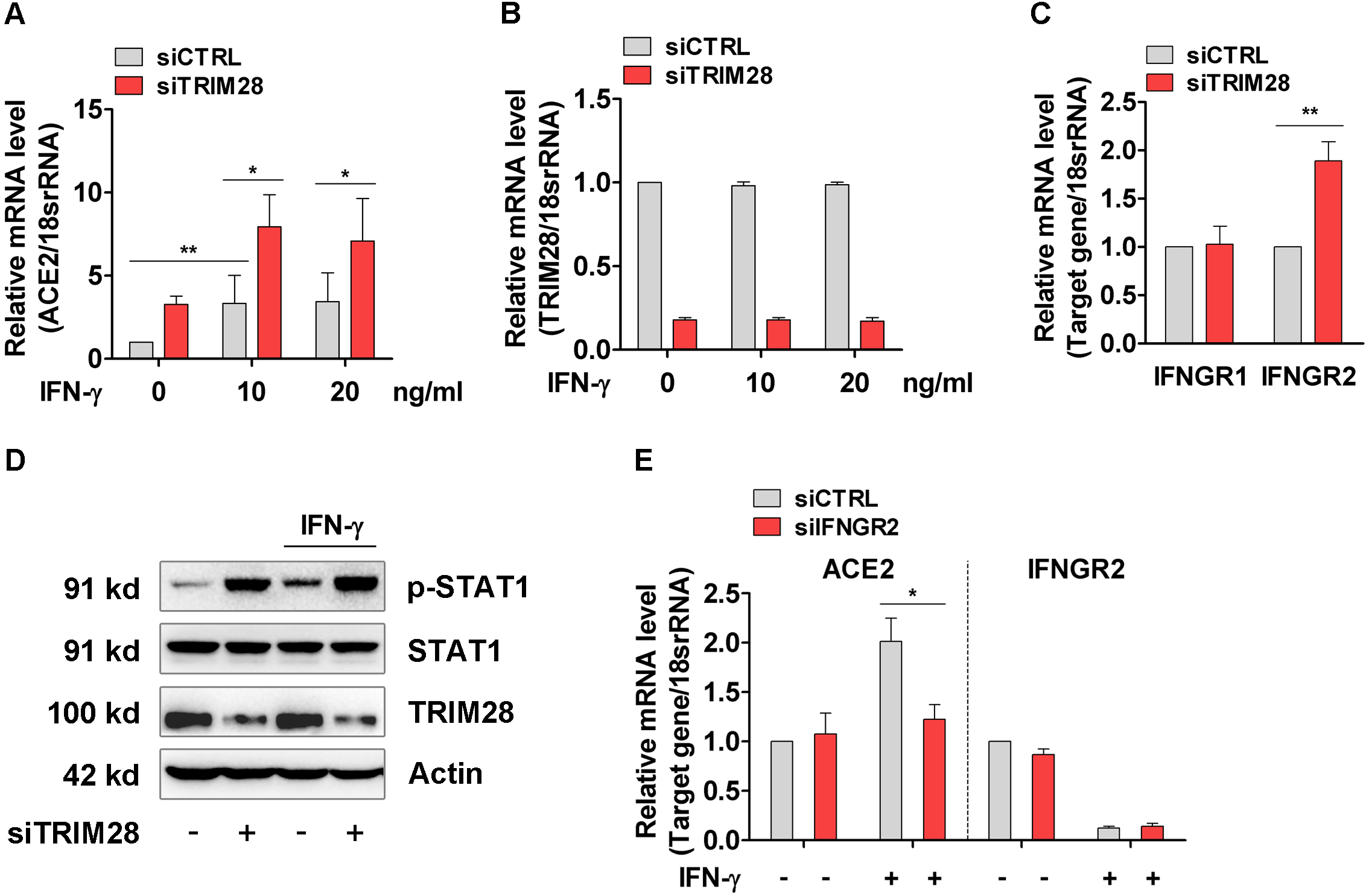
Effect of TRIM28 on IFN-γ induced ACE2 expression in A549 cells. (A, B) Control and TRIM28 silenced A549 cells were stimulated with IFN-γ for 16 h. qPCR assay was performed to check the expressions of ACE2 (A) and TRIM28 (B). n = 3. \*\**P*<0.01. \**P*<0.05. (C) The mRNA levels of IFNGR1 and IFNGR2 in control and TRIM28 knockdown A549 cells. n = 3. \*\**P*<0.01. (D) Control or TRIM28 knockdown A549 cells were treated with IFN-γ for 15 min. Immunoblot analysis was performed to detect expressions of phosphorylated STAT1 and total STAT1. Representative blots were shown. n = 3. (E) A549 cells were transfected with scramble or IFNGR2 siRNA, and then treated with IFN-γ (20 ng/mL) for 16 h. ACE2 and IFNGR2 expressions were examined by qPCR method. n = 3. \**P*<0.05.

### TRIM28 knockdown promotes ACE2 expression in human PAEpiCs

Finally, we validate the effects of TRIM28 on ACE2 expression in human PAEpiCs, a primary human lung epithelial cell. We observed that ACE2 was more abundantly expressed in TRIM28 knockdown PAEpiCs, compared to control PAEpiCs (Figure 5A). Consequently, pseudotyped SARS-CoV-2 cell entry was increased in human TRIM28 knockdown PAEpiCs, which can be reduced by ACE2 siRNA, indicating that TRIM28 knockdown promotes ACE2-dependent cell entry of SARS-CoV-2 (Figure 5B). Consistent with our mentioned results acquired in A549 cells (Figure 2), co-cultured with NK-92 cells also induced greater mRNA level of ACE2 in PAEpiCs, compare to control (Figure 5C). Considering that TRIM28 expression only slightly decreased whereas ACE2 expression dramatically increased in the PAEpiCs co-cultured with NK-92 cells, other mechanism regulating ACE2 expression maybe also exist. Finally, we evaluated the effect of TRIM28 knockdown on AEC2 in human PAEpiCs in the presence of dexamethasone and IFN-γ. Our results show that knockdown of TRIM28 still induces ACE2 expression of human PAEpiCs in the presence of dexamethasone. Meanwhile, knockdown of TRIM28 further enhanced the up-regulation of ACE2 in the presence of IFN-γ (Figure 5D, E). Furthermore, knockdown of TRIM28 was shown to increase the expression of IFNGR2 in PAEpiCs at basal level as well as in the presence of dexamethasone and IFN-γ (Figure 5F). These results support that, in human primary lung epithelial cells, knockdown of TRIM28 increased ACE2 expression and potentiated the inductive effect of IFN-γ on ACE2 through a mechanism involving upregulation of IFNGR2.

**Figure 5.**
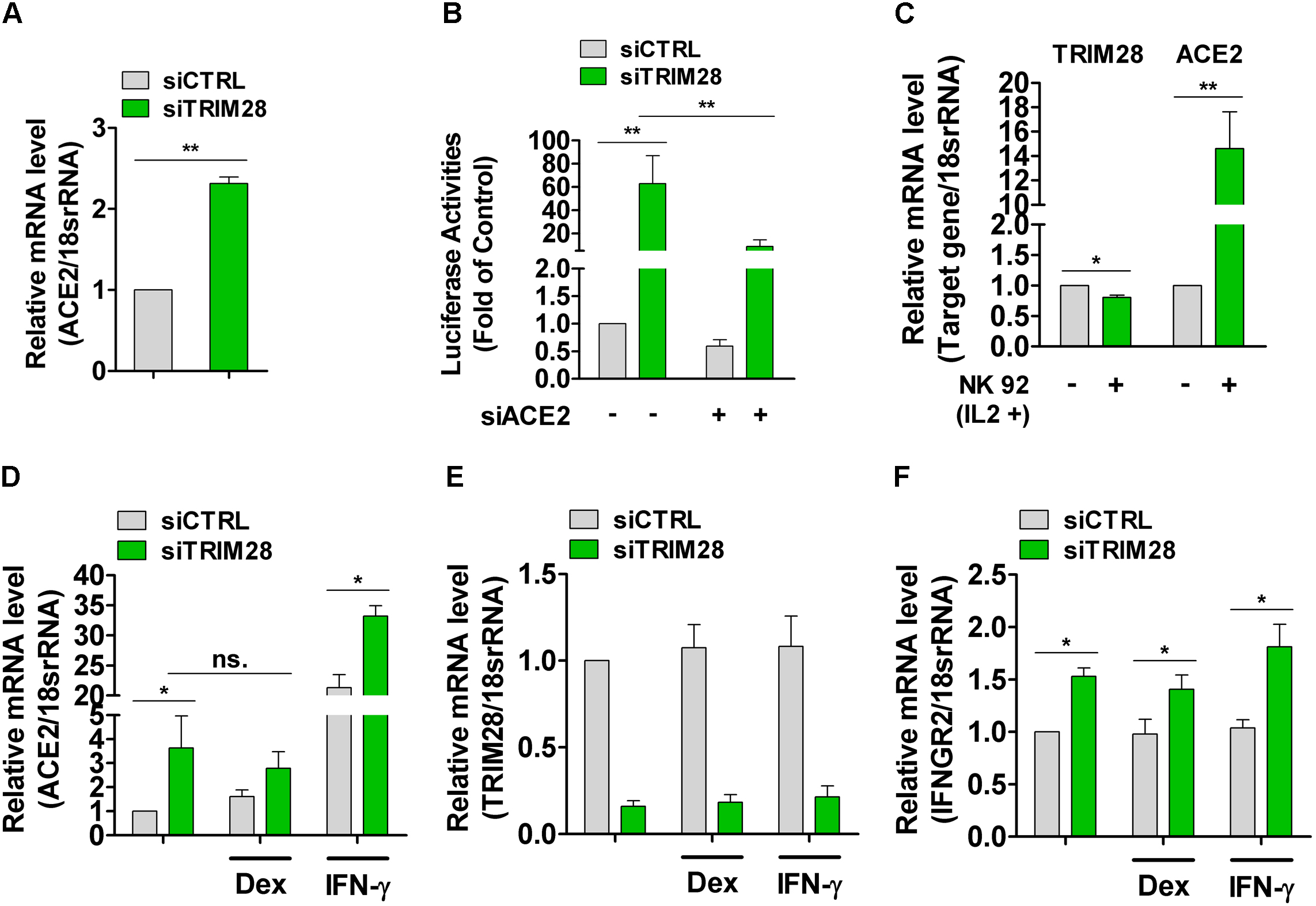
TRIM28 regulates ACE2 expression in human PAEpiCs. (A) ACE2 mRNA level in TRIM28 knockdown human PAEpiCs. n = 3. \*\**P*<0.01. (B) The human PAEpiCs were transfected with either scrambled or TRIM28 siRNA alone, or co-transfected with ACE2 siRNA for 24 h followed by infection with LV-Spike-nCoV-Luc. Forty-eight hours after infection, firefly luciferase activities were measured. n = 3. \*\**P*<0.01. (C) NK-92 cells were placed directly on top of human PAEpiC cell layer. After co-culture for 24 h, the mRNA expressions of target genes were analyzed by using qPCR method. n = 3. \*\**P*<0.01. \**P*<0.05. (D-F) Control and TRIM28 knockdown PAEpiCs were incubated with dexamethasone (1 μM) or IFN-γ (10 ng/ml) for 16 h, ACE2 (D), TRIM28 (E) and IFNGR2 (F) transcripts were analyzed by using qPCR assay. n = 3. \*\**P*<0.01.\**P*<0.05. ns. no significance.

## Discussion

In this study, we demonstrated that TRIM28 knockdown induces ACE2 expression in lung epithelial cells and increased pseudotyped SARS-CoV-2 cell entry. NK cell-derived factors inhibit TRIM28 and promote ACE2 expression in lung epithelial cells. Importantly, dexamethasone reduced the basal expression of ACE2 and partially reversed TRIM28 knockdown-induced upregulation of ACE2 in A549 cells. Furthermore, TRIM28 knockdown enhanced IFN-γ induced ACE2 expression through a mechanism involving upregulating IFNGR2 in both A549 and primary alveolar epithelial cells. Our study identified TRIM28 as a key regulator of ACE2 expression and SARS-CoV-2 cell entry and potential drug treatment targets for COVID-19.

Coronavirus entry is the first step of viral infection. The spike glycoproteins, localized on the envelope of coronaviruses, bind to their cellular receptors and then lead to the attachment and fusion between viral and host cell membranes [31–33]. Previous studies elucidated that ACE2 is engaged as SARS-CoV entry receptor. Cryo-electron microscopy revealed that one of receptor-binding domain of the trimeric spike glycoprotein can bind ACE2 and induce pre-to postfusion conformational transition [31, 34]. Recent studies reported that entry of SARS-CoV-2 in the host cell is also driven by binding of its spike protein with ACE2 and TMPRSS2 [1]. In normal human lung, ACE2 is mainly expressed in type II and type I alveolar epithelial cells. Single-cell RNA sequencing (scRNA-seq) data showed that ACE2 is expressed in 0.64% of all human lung cells, mainly in type II alveolar epithelial cells [35]. Furthermore, TRIM28 was reported to be co-expressed with ACE2 and TMPRSS2 among type II pneumocytes in a scRNA-seq dataset derived from surgical resections of fibrotic lung tissue [5]. We observed a relative high and low mRNA level for ACE2 in human PAEpiCs and A549 cells, respectively (data no shown), and showed that TRIM28 knockdown causes an increase in the expression of ACE2 and SARS-CoV-2 entry in both of two type cells. These data suggest that TRIM28 is a key regulatory factor for expression of ACE2, through which TRIM28 controls SARS-CoV-2 cell entry in lung epithelial cells

SARS-CoV-2 predominantly affects the lungs and can activate innate and adaptive immune responses and cytokine response in alveolar structures. scRNA-seq analysis showed an existing of myeloid dendritic cells, plasmacytoid dendritic cells, mast cells, NK cells, T cells and B cells in the bronchoalveolar lavage fluid from patients with severe COVID-19 [27]. Among these types of cells, NK cells functions to balance the immune responses by eliminating infected cells and oppressing dendritic cell and T cell activity, which are likely activated by SARS-CoV-2 infection and local cytokines in lung tissue. Moreover, the scRNA-seq analysis data support that NK cells can traffic into the lung where they contribute to local tissue injury, although lymphopenia was also observed in patients with severe COVID-19 [27, 28, 36]. In this study, we demonstrated that co-culture of NK cells with lung epithelial cells can decrease TRIM28 but increase ACE2 expression in lung epithelial cells. Our data highlight that when the initial virus infected-cells are eliminated by NK cells, the lung epithelial cells adjacent to NK cells, including virus infected- and non-infected cells, are more likely to become infected by SARS-CoV-2 due to enhanced expression of ACE2. Thus, the virus cell entry and NK cell infiltration can enhance each other, and form a positive-feedback loop to intensify lung inflammation and infection, suggesting NK cells acts as a double-edged sword against SARS-CoV-2, and may play a role in exacerbation of lung inflammation in patients with COVID-19.

Dexamethasone, a synthetic glucocorticoid compound with potent immunosuppressing activities, was recently found to reduce mortality of severe COVID-19 patients by one-third for patients on ventilators, and by one-fifth for those receiving oxygen [29]. This discovery has supported the usage of dexamethasone for management of COVID-19. However, the detail mechanism of how dexamethasone reduces mortality of severe COVID-19 patients is still unclear. Dexamethasone was shown to inhibit excessive cytokine production and prevent chemokine storm which damage pulmonary tissue. On the other hands, dexamethasone blocks naive T cell priming, expansion and differentiation, antibody making of T and B cells and activationof NK cells [37], and then potentially limit viral clearance. Moreover, patients with severe COVID-19 present lymphopenia showing that CD4+ T cell, CD8+ T cell, B cell and NK cell-count were reduced [27, 38]. Since that these immune cells contribute to the clearance of coronavirus and coronavirus infected cells, the timing of dexamethasone usage should be considered to avoid further reduction of lymphocytes [39]. In this study, we demonstrated here that dexamethasone inhibits expression of ACE2 of A549 cells at basal level as well upon in the presence of TRIM28 siRNA and NK cells, suggesting that decreased expression ACE2 is involved in the beneficial effect of dexamethasoneon restraining SARS-CoV-2. Meanwhile, we observed that dexamethasonedid not significantly alter the expresion of ACE2 in primary human PAEpiCs, highlighting that distinct strategy should be implanted for the effective prevention and treatment of the heterogeneous COVID-19 patients.

Previous studies have demonstrated a bidirectional crosstalk between IFNs and TRIM28. For example, IFN-α mediated immediate-early phase of transcriptional activation in HeLa cells was negatively regulated by TRIM28 [40]. The SUMOylation deficient TRIM28 can stimulate and prime IFN-α mediated innate immune defenses and restrict influenza A virus replication *via* a JAK1-dependent pathway [41]. Meanwhile, phosphorylation of TRIM28 leads to increased expression level of IFN-β in human lung epithelial cells during infection with highly pathogenic avian influenza viruses [42]. Knockdown of TRIM28 in A549 cells increased Sendai virus-induced IFN-α production and weakened IFN-sensitive vesicular stomatitis virus infection [30]. Furthermore, TRIM28 can bind with several IFN downstream effectors, and regulate IFN-dependent immune responses. For example, TRIM28 can bind with and increase SUMOylation of IRF7 during viral infections [30]. IRF5-mediated expression of TNF was inhibited by TRIM28 in human M1 macrophages [43]. TRIM28 associates with STAT1 and blocks IFN-induced STAT1-dependent IRF-1 gene expression [40]. The potential contribution of TRIM28 to IFN-γ stimulated ACE2 expression has been supported by the recent study which showed that TRIM28 is expressed in ACE2+/TMPRSS2+ type II pneumocytes [5]. We showed herein that TRIM28 knockdown can enhance expression of IFNGR2 and its downstream target gene, ACE2, in human lung epithelial cells, providing a new mechanistic link between TRIM28 and IFN-γ.

In summary, this study identified TRIM28 as a novel regulator of SARS-CoV-2 cell entry by regulating ACE2 expression in human lung epithelial cells. NK cells activation can inhibit TRIM28 but promote ACE2 expression in lung epithelial cells. Future studies focusing on the regulatory mechanism of TRIM28 expression will be of importance to understand the heterogeneity of COVID-19 patients and develop individualized preventive and therapeutic strategies. Our results also highlighted the potential distinct effect of dexamethasone on ACE2 expression in different COVID-19 patient groups, which may be further validated by the ongoing clinical trials of dexamethasone and the future studies.

## Acknowledgments

This study was supported by the National Natural Science Foundation of China (81870327, 81770443) and Clinical Advantage Discipline of Health System of Putuo District in Shanghai (#ptkwws201901 and #2019ysxk01).

## Author Contribution

PZ and YFW conceived and designed experiments. YFW, YZF, YTH, TD and PZ performed the experiments and analyzed the data. PZ, YW and NPW analyzed the data and drafted the manuscript. ZJL and DKH participated in the data analysis.

---

**Supplementary data 1.**
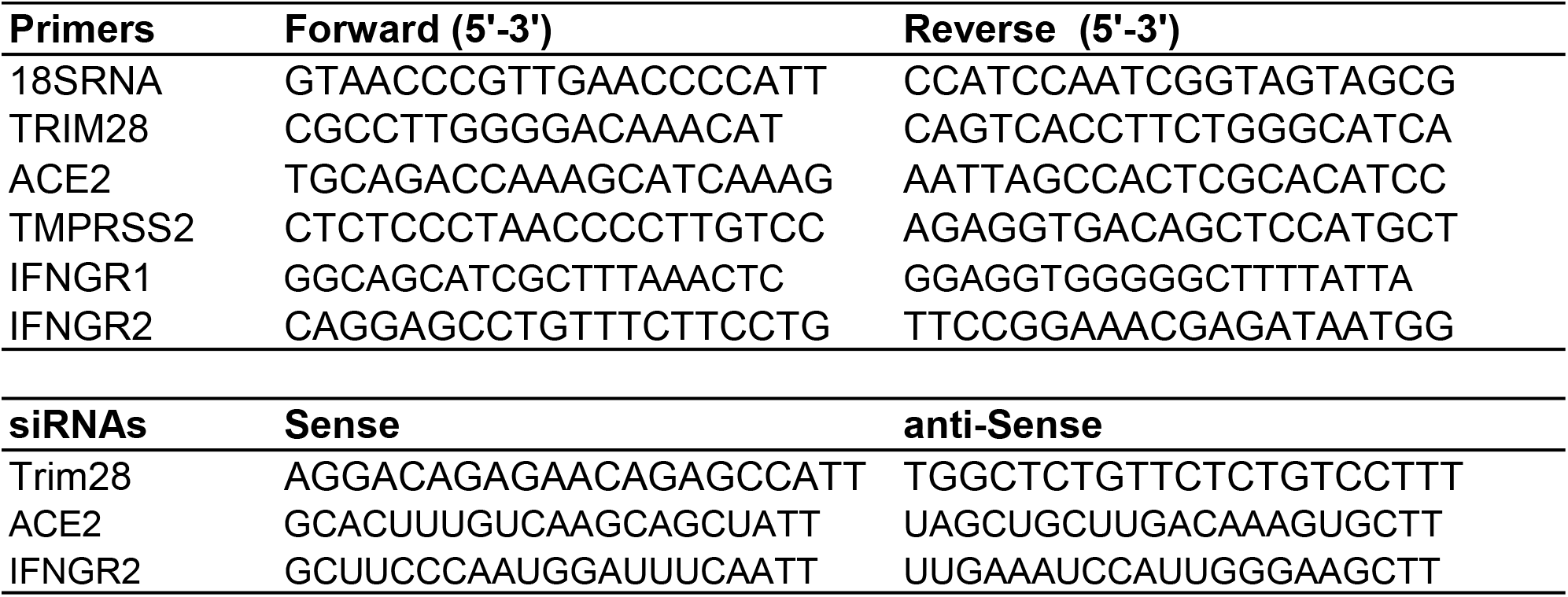
The primer and siRNA sequences used in this study.

